# A Theory of Heterosis

**DOI:** 10.1101/2025.02.24.639169

**Authors:** Zhao-Bang Zeng, Gabriel De Siqueira Gesteira, Lujia Mo, Yingjie Xiao, Jianbing Yan

**Author notes:** Corresponding author: Zhao-Bang Zeng, Bioinformatics Research Center, North Carolina State University, Raleigh, 27695-7566, NC, USA.

## Abstract

Heterosis refers to the superior performance of a hybrid over its parents. It is the basis for hybrid breeding particularly for maize and rice. Genetically it is due to interactions between alleles of quantitative trait loci (QTL) (dominance and epistasis). Despite enormous interest and efforts to study the genetic basis of heterosis, the relative contribution of dominance vs. epistasis to heterosis is still not clear. This is because most published studies estimate QTL effects in pieces, not able to put them together to assess the overall pattern adequately. We propose a theoretical framework that focuses on the inference of the relationship between genome and traits that includes the identification of multiple QTL and estimation of the whole set of QTL (additive, dominant, and epistatic) effects. Used for heterosis, it gives a clear genetic definition and interpretation of heterosis. We applied the theory and methods to a large maize dataset with a factorial design of many male and female inbred lines and their hybrid crosses. Heterosis of ear weight in maize is primarily due to QTL dominant effects, many are over-dominant. The contribution to heterosis due to epistasis is small and diffused. For comparison, we also analyzed a rice dataset that is an F2-type population derived from a cross between two inbred lines. The result indicates that dominance is still the main contributor to heterosis, and epistasis contribution is small.

**Article Summary:** We propose a general theoretical framework to analyze and interpret quantitative trait genetic variation in a population through the identification of quantitative trait loci (QTL) and the estimation of QTL effects including interactions. Applied to a large genomic study in maize, we produce direct estimation of genetic contribution to heterosis—QTL dominance and epistasis and compare them to the observed heterosis. The evidence is clear that the heterosis of ear weight in maize is primarily due to QTL dominance. The contribution to heterosis due to QTL epistasis is relatively small and diffused.

## Introduction

Heterosis refers to the superior performance of a hybrid over its parents. The utilization of heterosis is the basis of hybrid breeding, particularly in maize (Duvick 2005, Kusmec et al. 2021). The genetic basis of heterosis is due to the interaction of alleles of quantitative trait loci (QTL)—dominance and epistasis (*e*.*g*., Fujimoto et al. 2018). QTL Dominance has long been regarded as a major contributor to heterosis because it was thought that dominance can mask the effects of deleterious recessive alleles. Overdominance is an excessive form of dominance. There have been many reports of detection of QTL overdominance (Stuber et al. 1992, Krieger et al. 2010, Li et al. 2017), although cautions have been voiced that some may be due to pseudo-overdominance of multiple QTL in close repulsion linkage. Epistasis could also play a significant role in heterosis and there have been reports of detection of statistically significant epistasis between QTL alleles (*e*.*g*., Jiang et al. 2017). Despite tremendous interests and efforts, it is still largely unclear how to systematically study the genetics of heterosis and assess the relative importance of dominance (including overdominance) *vs*. epistasis on heterosis. One problem could be due to the limited sample sizes of many study populations which would limit our ability to assess QTL epistasis adequately. Another problem is that QTL effects including epistasis were typically estimated and explained in pieces in most studies (*e*.*g*., Huang et al. 2016; Xiao et al. 2021) which would impose the difficulty to assess the overall effects consistently and in totality.

In this study, we propose a general multiple QTL genetic model to model the relationship from genome to phenotypes in a population for general quantitative genetics data analysis and interpretation and use it to study heterosis. This approach could produce a direct estimate of the genetic composition of heterosis, thus providing evidence to assess the relative importance of dominance *vs*. epistasis on heterosis and many genetic questions in the study population.

### Theory and Methods

Although this theory is currently proposed for the study of heterosis, the theoretical framework can be used for many applications and studies of quantitative trait variation in complex populations and environments, particularly for the evolutionary study of complex traits in general. This approach would work better for a large population with dense genetic markers.

Traditionally the study of heterosis uses a cross (F1) of two inbred lines (P1 and P2) and from F1 to produce an F2 population or further crosses. This design studies the heterosis between an F1 and the mean of P1 and P2 and uses F2 or other segregating populations to detect and estimate QTL effects and relate those estimates to the observed heterosis. An example is the study of Hua et al. (2002, 2003) on rice heterosis.

Recently, Xiao et al. (2021) reported a study of maize heterosis, that has 6210 crosses (F1’s) between 30 male inbred lines (P1’s) from one heterotic group and 207 female inbred lines (P2’s, drawn from 1404 recombinant inbred lines resulted from multiple-way crosses of 24 original lines) from another heterotic group, with extensive genomic genotypes and trait phenotypes. This is a North Carolina Design II (factorial design) population. In this population, each F1 hybrid has different parents and different heterosis. We will use these two studies to lay out our theory and discuss the genetic basis of heterosis in maize and rice. Of course, the theory can be used or adapted for other experimental designs and data structures. We first focus on the study of Xiao et al. (2021)

### Population

Let *P*1_*i*_ for *i* = 1,2, …, *n*_1_ and *P*2_*j*_ for *j* = 1,2, …, *n*_2_ be inbred lines of the two heterotic groups and *F*_*ij*_ be their hybrids. We treat inbred lines and their hybrids as one population for model specifications, genetic estimation, and interpretation.

### G2A genetic model

Suppose we observe genotypes of many genomic SNP markers for *P*1_*i*_ and *P*2_*j*_ and hence *F*_*ij*_ (deduced from the marker genotypes of *P*1_*i*_ and *P*2_*j*_). If the marker coverage is dense, we can practically treat some markers as potential QTL and perform marker selection analysis. As SNP makers are biallelic, we will treat potential QTL as biallelic. If some markers of *P*1_*i*_ and *P*2_*j*_ are heterozygous, those markers can be accommodated in the analysis or treated as missing data.

We will use a general two-allele (G2A) model (Zeng et al. (2005) and Wang and Zeng (2006)) to model and analyze QTL in both inbred lines and hybrids. A G2A model, rather than an F2 model, is more appropriate here to describe the genetic variation of the population, as allelic frequencies of markers and potential QTL are very uneven *among the male and female inbred lines (the source of variation)*.

What is the G2A model? We first give the G2A model a general derivation and explanation. Let *s* ∈ (*P*1_*i*_, *P*2_*j*_, *F*_*ij*_; *i* = 1,2, …, *n*_1_, *j* = 1,2, …, *n*_2_). We use *z*_*slg*_ to index the allelic state of individual *s*, locus *l* (*l* = 1,2,…,*m*) and gamete *g* (*g* = 1,2). We consider two alleles *A*_*l*1_ and *A*_*l*2_ that are segregating in the population with allele frequency *p*_*l*_ for *A*_*l*1_. Let

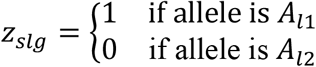

We use a centralized variable *x*_*slg*_ = *z*_*slg*_ − *Prob*(*z*_*slg*_ = 1) = *z*_*slg*_ − *p*_*l*_ for model setting.

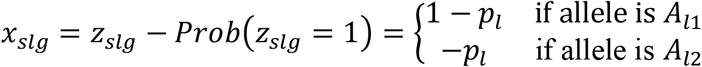

Why centralizing variables? This is a well-known statistical practice. Centralizing or standardizing variables is especially important when a regression model contains interaction terms. If variables are not centralized or standardized when a model contains these types of terms, there is a risk of missing statistically significant results or producing potentially conflicting results, *i*.*e*., the model is internally inconsistent. The consistency means that a lower-dimension model is consistent in a higher-dimension space *under certain conditions* (Zeng et al. 2005).

With this specification, the relationship between a trait phenotype *y*_*s*_ and genotypes of multiple QTL (*x*_*slg*_′*s*) can be modeled as

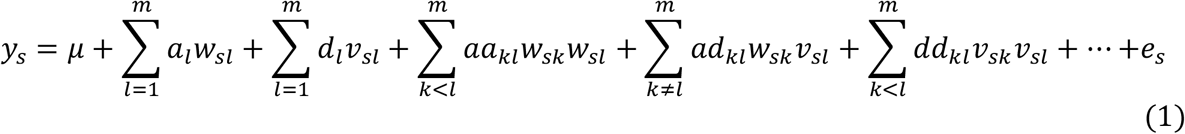

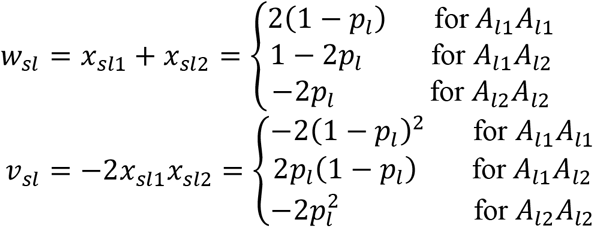

with 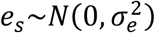. In this model, *w*_*sl*_ is additive allelic variable and *a*_*l*_ is additive effect of QTL *l, v*_*sl*_ is dominant (additive-by-additive interaction) variable and *d*_*l*_ is dominant effect of QTL *l, aa*_*kl*_, *ad*_*kl*_, and *dd*_*kl*_ are additive-by-additive, additive-by-dominant, and dominant-by-dominant epistatic effects between QTL *k* and *l* respectively.

We used the G2A model form of Zeng et al. (2005) with *v*_*sl*_ = −2*x*_*sl*1_*x*_*sl*2_, differing on the specification of *v*_*sl*_ = *x*_*sl*1_*x*_*sl*2_ of Wang and Zeng (2006) by a factor −2. This is for the purpose to be in line with the specification of the F2 model (symmetric model) with *p*_*l*_ = 1/2. The F2 model was first introduced by Anderson and Kempthorne (1954) and has been used extensively in literature for many applications.

Why G2A model for general applications? This G2A model is a digital model or binary model that has simplicity in setting and can represent any complexity in multitude. Biologically, since many genomic studies contain dense SNP markers that are bi-allelic, it is reasonable to assume that some SNP markers are targeted QTL or very closely linked to casual variants, and a model selection from those SNP markers can have a good representation of the genetic structure of quantitative trait variation in a population. Theoretically, this model is akin to the infinite site mutation model (Crow and Kimura 1970). How about multiple alleles? The case of multiple alleles can be conveniently represented by multiple two-alleles (SNP).

Incidentally, this G2A model can be readily extended to polyploids. Let *ρ* be ploidy number (2 for diploid, 4 for tetraploid, 6 for hexaploid, etc). In the model above, we can extend the gametic index to *ρ* (*g* = 1,2,…,*ρ*). Then, the allelic dosage additive variable can be extended to 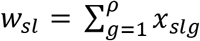. The summation is the concept of allelic dosage for diploid and polyploids. Correspondingly, the pair dosage (two allelic interaction) dominant variable 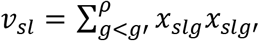, the triplet dosage (three allelic interaction) dominant variable 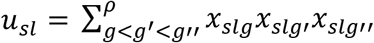, etc. With these specifications, a polyploid G2A model can be expressed in a similar form of equation (1) for all levels of allelic effects and interactions within and between loci. A more general discussion on the implications and applications of this polyploid G2A model will be presented elsewhere.

### Heterosis

Heterosis of a trait for *F*_*ij*_ from *P*1_*i*_ and *P*2_*j*_ is defined as

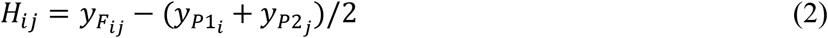

To study the genetic basis of heterosis, we need to show what constitutes heterosis in genetic terms. Denote 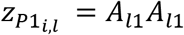 or *A*_*l*2_*A*_*l*2_ for *l* = 1, 2, …, *m* and similarly for 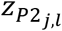 as well for marker genotypes of inbred lines. Then

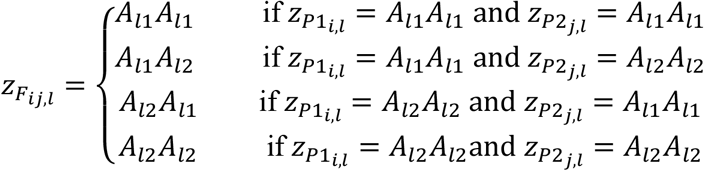

By using the above genetic model, the heterosis is expected to be as follows if we restrict the analysis to additive, dominant and additive-by-additive epistatic effects.

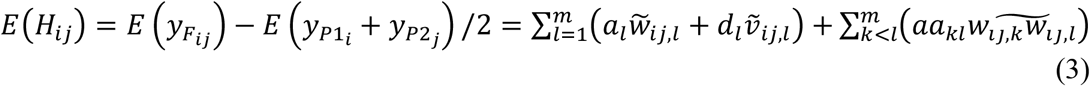

with

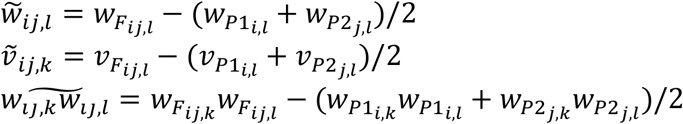

For a QTL locus if both parents have the same homozygote (either both *A*_*l*1_*A*_*l*1_ or *A*_*l*2_*A*_*l*2_), the hybrid genotype is still *A*_*l*1_*A*_*l*1_ or 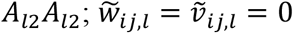. When parental genotypes have different homozygotes *(i*.*e*., one is *A*_*l*1_*A*_*l*1_ and the other is *A*_*l*2_*A*_*l*2_), the hybrid is heterozygote (*A*_*l*1_*A*_*l*2_) and

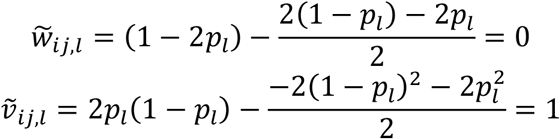

Thus,

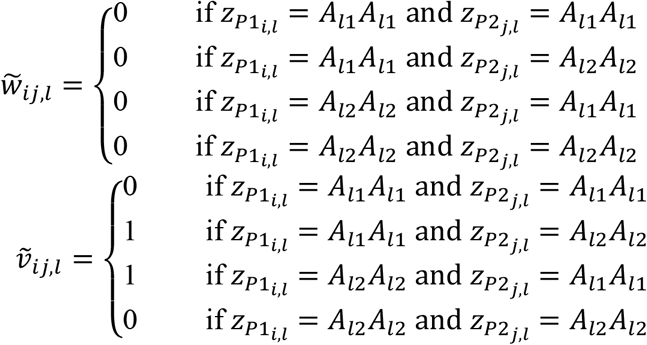

This shows that *a* does not contribute to heterosis and *d* contributes to heterosis for loci in heterozygote in hybrid.

For a pair of QTL (*k* and *l*), if 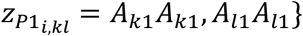 and 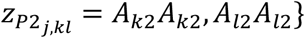 or 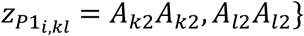 and 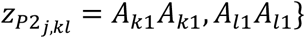

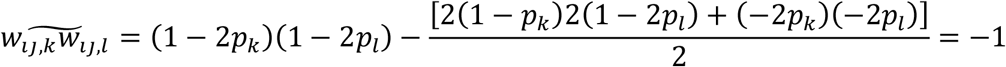

If 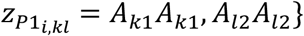 and 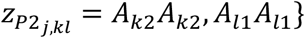 or 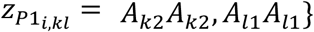 and 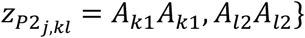

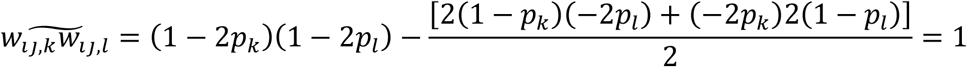

For all other cases, 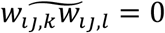. Thus,

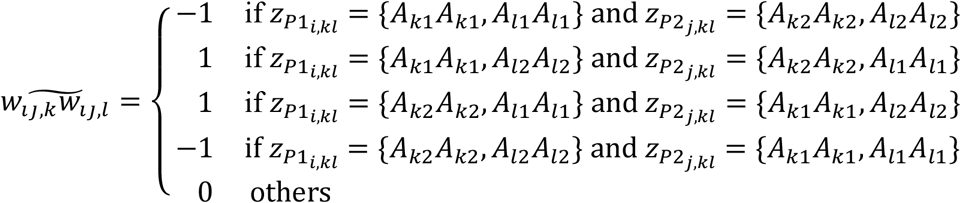

This shows that *aa* contributes to heterosis if both parents have different homozygotes on the two loci.

Thus, given estimates of genetic model parameters (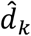 and 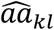), we can estimate genetic partitions and components of heterosis for each hybrid.

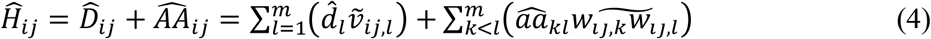

This can be compared with the observed heterosis *H*_*ij*_.

Here we ignored additive-by-dominant 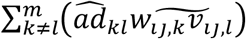, dominant-by-dominant 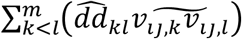 and higher order epistatic effects because they are higher-order statistics and less important. The terms 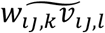 and 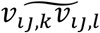 are complex and non-zero in general but are zero when allele frequency *p*_*k*_ = *p*_*l*_ = 1/2. That is, for the F2 model with allele frequency half, additive-by-dominant and dominant-by-dominant epistasis does not contribute to heterosis (Melchinger et al. 2007, Garcia et al. 2008). If deemed necessary, we could include additive-by-dominant and dominant-by-dominant epistasis in the analysis.

Note that, as pointed out in Garcia et al. (2008), the interpretation of heterosis genetics depends on the genetic model used. A commonly used genetic model in quantitative genetics literature is the *F*_∞_ genetic model largely because of its simplicity in expression. Based on the *F*_∞_ model, heterosis also depends on dominant-by-dominant interaction. In this study, we used a G2A model and its special form *F*_2_ model for the reason explained in Zeng et al. (2005). It is based on the principle of partition of genetic variances, the legacy of Fisher (1918).

Now we summarize and highlight a few points of our model-based analysis approach. First, we put inbred lines and hybrids together as one population and model the genetic variation for the whole population. As such, a model selection and estimation can explain the genetic variation within and between inbred lines and hybrids. The heterosis is defined specifically for each pair of inbred lines and their hybrid and is shown to be due to dominant effects of QTL that are heterozygote in the hybrid and *aa* epistatic effects of QTL that have opposite homozygote genotypes of the two loci in the inbred lines. In this model system, we try to identify individual QTL and analyze additive effects of QTL alleles and significant pair-wise interaction effects of QTL alleles (dominant effects within loci and *aa* epistatic effects between loci). Heterosis is all about the interactions of QTL alleles, primarily the pair-wise allelic interaction effects of QTL (dominance within loci and *aa* epistasis between loci).

## Data and analysis

We applied this model to the dataset of Xiao et al. (2021). The data consists of the high-quality whole genome SNP marker (~4.5 million) genotypes of 1428 inbred lines from the CUBIC (Complete-diallel plus Unbalanced Breeding-derived Inter-Cross) synthetic population as a maternal pool and 30 paternal tester lines from diverse genetic backgrounds. We will focus our analysis and discussion on a cross population that includes 207 maternal lines (randomly selected from the 1428 CUBIC lines) and 30 paternal lines and their 6210 hybrids. Twenty quantitative traits were measured in five locations for all inbred lines and hybrids. Our analysis is primarily focused on the inference of genetic structure of the population on each quantitative trait, and through the inference to study the genetic basis of heterosis.

### Model selection

Model selection is at the core of a model-based analysis. The results will depend on the selection procedures and criteria. We used LASSO method and combined it with stability selection for model selection. LASSO (Tibshirani 1996) is a statistical method that shrinks regression coefficient estimates through the L1 regularization, leading many small estimates to zero to achieve a subset selection. Stability selection ((Meinshausen and Bühlmann 2010) uses LASSO model selection in sub-sampling data to explore model structure. It provides an algorithm for selecting a model while controlling the number of false discoveries.

Specifically, this is the procedure we used for data analysis. First, there are too many markers in the data for marker subset selection analysis. To achieve a feasible and efficient computation, we initially generated an evenly spaced marker subset by sampling one marker every 800 markers across the whole genomes of 10 chromosomes. After removing markers with minor allele frequency (MAF) less than 1/(30+24)=0.0185 and with genotype errors and contamination, a total of 4,701 markers were retained in the marker pool for subsequent model selection. We used stability selection (Meinshausen and Bühlmann 2010) to enhance the consistency and robustness of QTL selection while controlling false discoveries. The procedure involved 100 times of random subsampling from the total 6,447 samples, with each round drawing 50% of the samples without replacement. Marker selection for each of the 100 sample sets was conducted using group LASSO (Simon et al. 2013) via the R package ‘grplasso’ (Meier et al. 2008) along with 10-fold cross-validation, in which the additive and dominance effects of a QTL were selected together as a pair. Given the tunning parameter *λ* with the minimum cross-validation error, QTL consistently selected in at least 50 of the 100 subsamples were retained for further analysis. Conditional on the selected additive and dominant QTL effects, the same selection procedure was used again to identify additive-by-additive epistatic interactions among all combinations of those selected QTL. The QTL (*a, d*, and *aa*) effects in the final selected model were re-estimated in the full sample.

Thus, we used a two-step selection procedure, first selecting QTL main effects (putting *a* and *d* together in selection) from candidate markers, then selecting QTL *aa* effects only from the QTL pairs selected in the first step. This procedure was in line with a similar procedure used in Laurie et al (2014) which explained the justification and rational for the multiple step selection procedure.

During the investigation, we selected and compared many different models. A detailed discussion on model comparison is complex. Although some model details may vary for different model selection procedures, the genetic result pattern and conclusions reported below are robust. To simplify the result report and discussion, we report the results based on a representative selected model.

## Results and Discussion

Given an inferred genetic model for a study population, one can explore the genetic structure of the population for a quantitative trait, estimate or predict any quantity including its genetic components, trace the causes (QTL changes) and process (a sequence of changes) of evolutionary events or selection responses to breeding efforts, and utilize the inferred information for a more proactive or creative intervention (*e*.*g*., breeding design). These are just a few examples of advantages and opportunities that a model-based quantitative genetic inference can and should play for an agricultural breeding program or any biological inquiry on complex traits.

### Selected QTL model for ear weight and heterosis composition

Since ear weight is the most important trait economically and has the most significant heterosis, we will discuss the model selection results mostly using ear weight as an example and then discuss other traits for comparisons. Based on the selection procedure, we selected a model of 139 QTL distributed all over the genome with additive (*a*) and dominant (*d*) effects and selected 413 additive-by-additive (*aa*) epistatic effects (supplemental Table S1).

Figure 1 plots some details of this inferred genetic model of ear weight: genotypes (A_1_A_1_ or A_2_A_2_), estimated additive (*a*) and dominant (*d*) effects (red and blue dots) of 139 QTL for 30 male lines, 24 female founder lines, and 183 derived female lines. This figure provides a visual picture of the genetic differences between the two heterotic groups (male and female lines) on ear weight. It shows what contributes to the heterosis and how one might design a better breeding plan to further improve the breeding lines.

**Figure 1:**
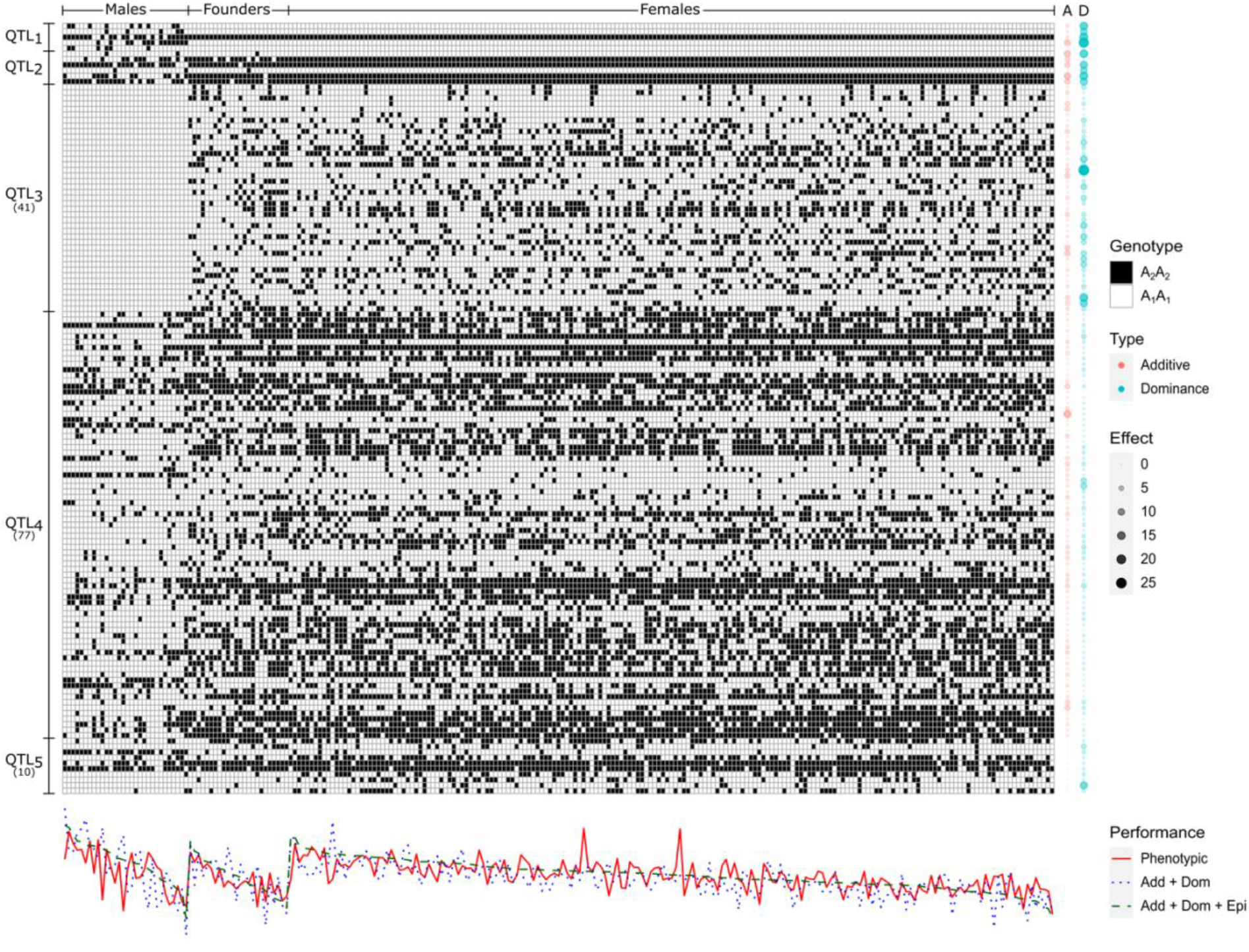
The inferred genetic model of ear weight: genotypes (*A*_1_*A*_1_ or *A*_2_*A*_2_), estimated additive (*a*) and dominant effects (*d*) (red and blue dots grouped in absolute size) of 139 QTL for 30 male lines, 24 female founder lines and 183 derived female lines arranged in columns. QTL are arranged in rows and grouped to show the pattern of the genetic structure in the parental populations. Group 1: five QTL that segregate only in males; Group 2: six QTL that segregate only in males and founders; Group 3: 41 QTL that segregate in founders and derived females, but not in males; Group 4: 77 QTL that segregate in males, founders and derived females; Group 5: ten QTL with additive effects close to zero, but significant dominant effects. The lower panel shows observed phenotypic values of ear weight, estimated ear weights with QTL additive and dominant effects, and with QTL additive, dominant, and additive-by-additive effects.

More genetic details are plotted in Figure 2. Figure 2A plots the distribution of 139 QTL with additive (*a*) and dominant (*d*) effects and 413 additive-by-additive (*aa*) epistatic effects on ear weight. A distinct feature emerges that *a* and *aa* effects are centered around zero, about equally positive or negative, and *d* effects are predominantly positive. This goes right in the heart of the genetic interpretation of the causes of heterosis. As seen clearly in Figure 2E which shows the estimates of heterosis components (dominance *vs. aa* epistasis, equation (4)), the conclusion is clear that the heterosis in ear weight is primarily due to QTL dominant effects. The contribution due to epistasis is small in magnitude and non-directional. Further, by Figure 2B which plots dominant (*d*) effects of QTL against the degree of dominance (*d/*|*a*|), QTL effects on heterosis on ear weight are overwhelmingly over-dominant. It is possible that some of those over-dominant QTL could be pseudo-overdominance due to multiple underlying genes in close repulsion linkage. But the evidence on over-dominance is overwhelming.

**Figure 2:**
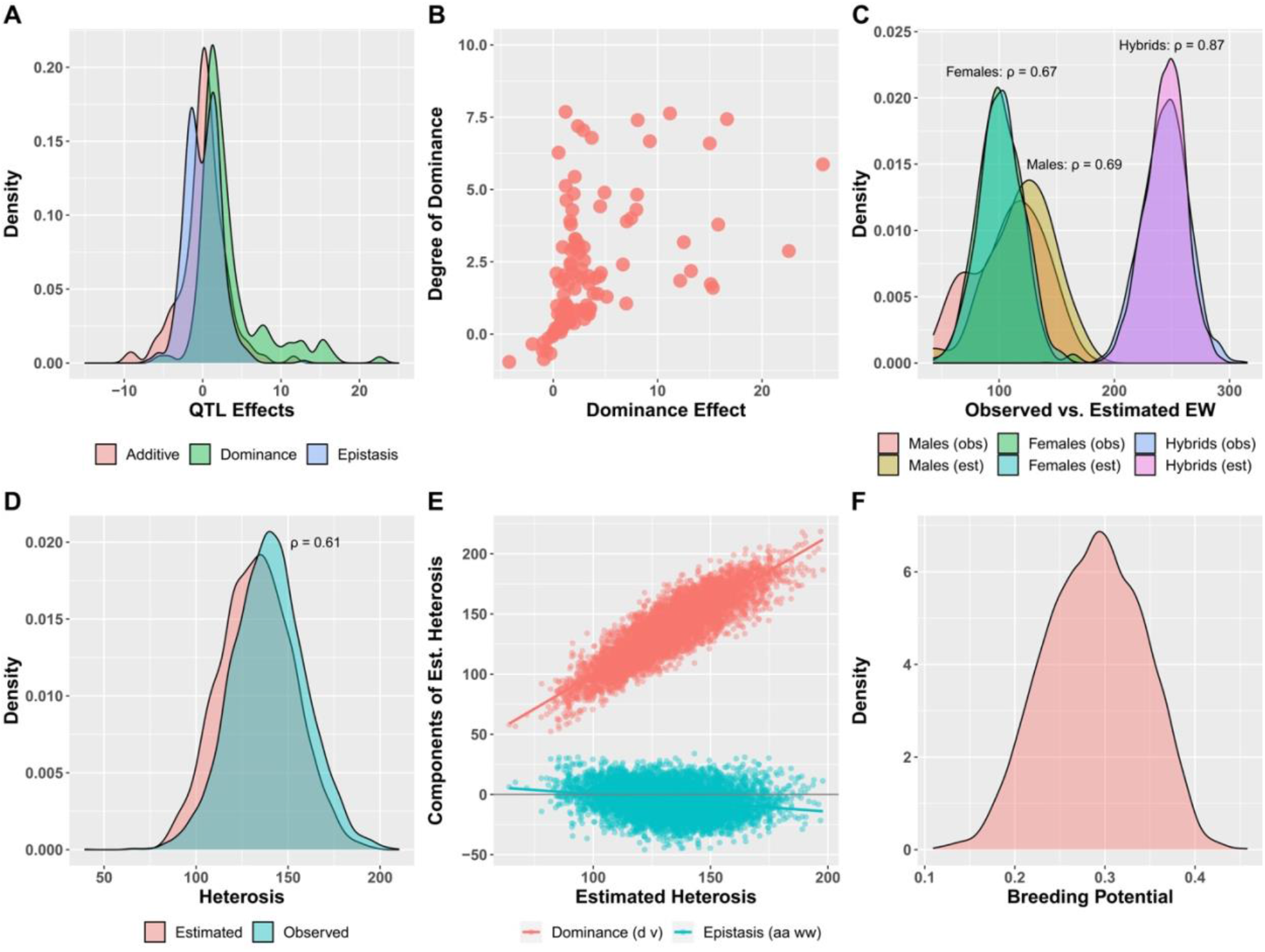
Estimation of genetic model and associated properties on ear weight in the population. **A:** The distribution of 139 QTL with additive (*a*) and dominant (*d*) effects and 413 additive-by-additive (*aa*) epistatic effects on ear weight. **B**: Dominant (*d*) effects of QTL are plotted against the degree of dominance (*d/*|*a*|). **C**: The comparison of estimated *vs*. observed ear weights for (male and female) inbred lines and hybrids. **D**: The comparison of estimated *vs*. observed ear weights for heterosis. **E**: The distribution of the components (dominance *vs. aa* epistasis, equation (4)) of estimated heterosis for 6210 hybrids is plotted against the estimated heterosis. **F**: This plots what may be called the breeding potential of 6210 hybrid crosses: the dominant contribution 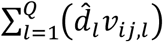 of each hybrid to heterosis as a ratio of the maximal contribution of QTL dominant effects to heterosis 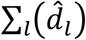, if all segregating QTL alleles are in heterozygote in a hybrid, *i*.*e*., 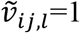 for all *l*.

Figures 2C and 2D show the model fit, the comparison of estimated *vs*. observed ear weights for inbred lines and hybrids, and their differences (heterosis). More details are provided in Table 1 on the partition of variances and covariances of genetic components (A, D, and AA) and residuals in male inbred lines, female inbred lines, hybrids, and heterosis (the difference between hybrid and mean of inbred parental lines) and the broad-sense heritability. For male inbred lines, female inbred lines, and hybrids, the intercept (*μ*), A, D, AA, and residual are specified by equation (1), and for heterosis, D, AA, and residual are specified by equations (2) and (4).

**Table 1:** Partition of variances and covariances of genetic components (A, D, and AA) and residuals in male inbred lines, female inbred lines, hybrids, and heterosis (difference between hybrid and mean of male and female parental lines) and the broad-sense heritability.

|  | Intercept | Partition of Variance |  |  |  | Heritability |
| --- | --- | --- | --- | --- | --- | --- |
| Intercept | 218.86 | A | D | AA | Residuals | H<sup>2</sup> |
| Male | A | 355.60 |  |  |  | 0.71 |
|  | D | 273.78 | 804.59 |  |  |  |
|  | AA | -261.72 | -522.73 | 635.98 |  |  |
|  | Residuals |  |  |  | 519.33 |  |
| Female | A | 448.20 |  |  |  | 0.77 |
|  | D | -215.04 | 486.45 |  |  |  |
|  | AA | -103.86 | -101.73 | 268.73 |  |  |
|  | Residuals |  |  |  | 237.06 |  |
| Hybrid | A | 195.45 |  |  |  | 0.79 |
|  | D | -56.10 | 181.60 |  |  |  |
|  | AA | -20.50 | 11.67 | 59.70 |  |  |
|  | Residuals |  |  |  | 97.95 |  |
| Heterosis | D |  | 674.70 |  |  | 0.67 |
|  | AA |  | -222.35 | 165.45 |  |  |
|  | Residuals |  |  |  | 308.44 |  |

Figures 2C and 2D and Table 1 show the fit of the genetic model to the population on ear weight in different ways.

The model fit for female parents and hybrids is better than that for male parents. This is partly because there are more female parents (207) and hybrids (6210) than male parents (30) that differentially contributed to the overall model fit, but probably more because the female parents have their genomes randomized during the construction of CUBIC lines and the male parents are genetically more diverse. It is notable that hybrids have smaller residual variances (Table 1) than their parents, which is generally observed in hybrid experiments.

### Degree of heterosis

In this study, each hybrid has different parents and thus different heterosis. The degree of heterosis varies from hybrid to hybrid and from trait to trait. We define the degree of heterosis by

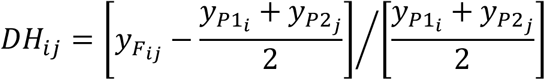

which is plotted in Figure 3A. There is a big difference in DH between yield-related traits, such as EW (ear weight) and KWPE (kernel weight per ear), and other traits. This reflects the fact that the selection for heterosis has been primarily concentrated on yield in the past hybrid breeding in maize and resulted in a profound degree of heterosis. Genetically, the difference in DH between traits is related to the mean degree of dominance 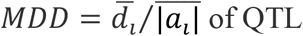. Figure 3B plots the mean degree of heterosis (MDH) against the mean degree of dominance (MDD). This means that selection on heterosis can increase the degree of dominance on QTL.

**Figure 3:**
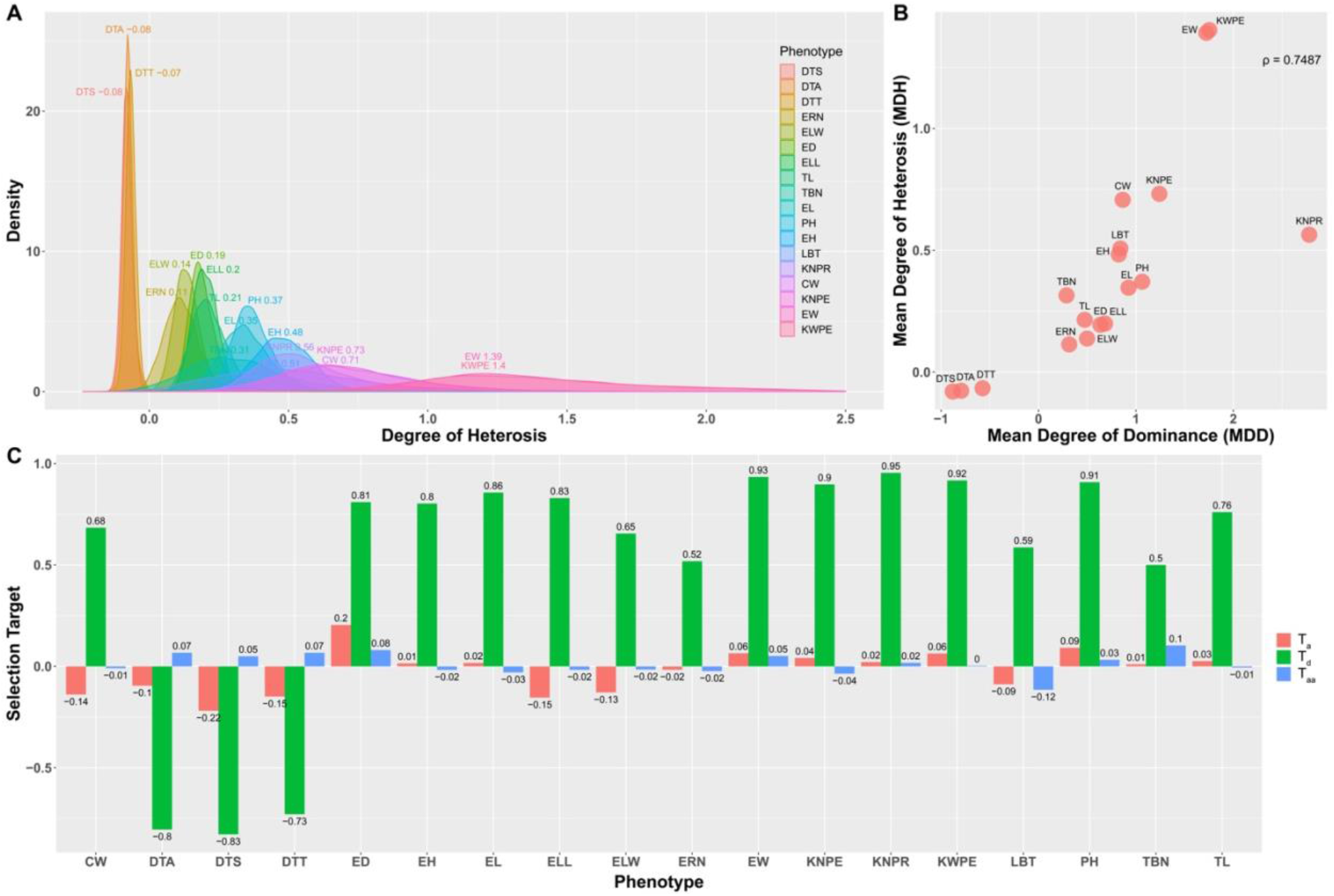
**A**: The degree of heterosis of 6210 hybrids is plotted for the 18 traits measured in the experiment. **B**: The mean degree of QTL dominant effects inferred for the 18 traits are plotted against the mean degree of heterosis for the 18 traits. **C**: Selection target of QTL effects (*T*_*a*_, *T*_*d*_, and *T*_*aa*_) inferred for the 18 traits. Name of the traits: CW (cob weight), DTA (days to anthesis), DTS (days to silking), DTT (days to tasselling), ED (ear diameter), EH (ear height), EL (ear length), ELL (ear leaf length), ELW (ear leaf width), ERN (ear row number), EW (ear weight), KNPE (kernel number per ear), KNPR (kernel number per row), KWPE (kernel weight per ear), LBT (length of the barren tip), PH (plant height), TBN (tassel branch number), TL (tassel length).

### Selection target

The distribution of estimated QTL effects in Figure 2A immediately suggests a measure that measures the direction of QTL effects, which may be called selection target as it tends to reflect the impact of selection: 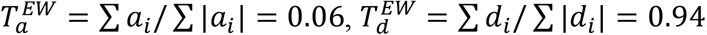, and 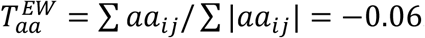. Figure 3C shows the measures for all traits in the study.

This is of course due to hybrid breeding that has been practiced for over a century (Duvick 2005, Kusmec et al. 2021). Hybrid breeding explored heterosis between different heterotic groups by using reciprocal recurrent selection that is targeted to preserve and enhance heterosis. As a result, hybrid performance and heterosis have been consistently enhanced over time, although surely inbred lines have also been continuously improved particularly with significant efforts to eliminate deleterious recessive alleles that hinder inbreeding efforts. Thus, the primary selection targets are QTL dominant effects. Male and female inbred line performance is similar for ear weight and other traits; thus, QTL additive effects are not directional. Additive-by-additive epistatic effects are also not directional for the population. This, of course, agrees with the conclusion that the QTL additive-by-additive epistatic effects are not the main cause and contributor to the maize heterosis as shown in Figure 2E.

This QTL effect-based measure is very useful to detect and explain the cause and/or consequence of evolutionary process. In a large-scale QTL mapping study between *Drosophila simulans* and *Drosophila mauritiana* on male genital arch (MGA), 19 QTL were detected with estimates of QTL effects in two backcrosses (Zeng et al. 2000). From Table 2 of Zeng et al. (2000), we can take the averages of QTL effects in two backcrosses as the estimates of QTL additive effects. This gives 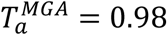 with 18 of the 19 QTL having additive effects in one direction. We can also take the differences of QTL effects in two backcrosses as the estimates of QTL dominant effects. This gives 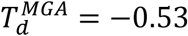. This gives strong evidence that the male genital arch may have been strongly selected in opposite directions during and since speciation due to differential female preferences in the two species (True et al. 1997) and the impact of directional selection is primarily reflected on QTL additive effects.

**Table 2:** Observed and estimated heterosis and parental line differences for the four traits in the rice heterosis study. The estimated heterosis includes the components due to QTL dominance and epistasis and their respective percentages. Estimated mean degree of dominance 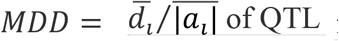 for the four traits.

|  |  | Yield | Tillers | Grains | Grain weight |
| --- | --- | --- | --- | --- | --- |
| **Heterosis** | Observed | 28.9 | 3.4 | 55.05 | 2.48 |
|  | Estimated | 33.35 | 8.18 | 49.12 | 1.26 |
| **Parental Differences** | Observed | -23.03 | -3 | -43.5 | -5.95 |
|  | Estimated | -10.33 | -1.77 | -25.62 | -3.68 |
| $\sum d$ | | 32.55 | 7.78 | 23.45 | 0.96 |
| $\sum aa$ | | -0.79 | -0.4 | -25.67 | -0.3 |
| % of $\sum d$ | | 0.98 | 0.95 | 0.48 | 0.76 |
| % of $\sum aa$ | | 0.02 | 0.05 | 0.52 | 0.24 |
| Mean degree of dominance |  | 2.86 | 2.54 | 1.69 | 0.56 |

This conclusion is mirrored in a large-scale directional artificial selection experiment in *Drosophila melanogaster* on wing shape (WS) (Weber 1990, 1992). After intensive bi-directional selection on wing shape for 16 generations, a high line and a low line were created and then crossed to create segregating recombinant inbred lines for QTL analysis. For chromosome 3, 11 QTL were detected with 10 QTL additive effects in one direction (Table 5 of Weber et al. (1999)). This gives 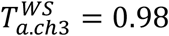. QTL analysis also detected 9 additive-by-additive epistatic effects with 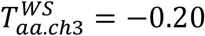. For chromosome 2, this QTL pattern was also repeated with 10 QTL detected and all additive effects in one direction, thus 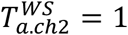 and 14 additive-by-additive epistatic effects detected with 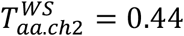 (from Table 3 of Weber et al. (2001)).

When we put all these together, we can conclude that a QTL model-based analysis can reveal a selection target that reflects the action and consequence of the selection process. For directional selection, it is reflected in QTL additive effects, whether it is natural directional selection as in the case of *D. simulans* and *D. mauritiana* on male genital arch or artificial directional selection as in the case of *D. melanogaster* on wing shape. For hybrid breeding, it is reflected in QTL dominant effects.

### Breeding potential

Since QTL dominant effects are the primary contributor to heterosis, it is of interest to examine the dominant contribution 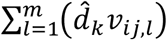 of each hybrid to heterosis as a ratio of the maximal contribution of QTL dominant effects to heterosis 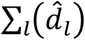, if all segregating QTL alleles are in heterozygote in a hybrid, *i*.*e*., 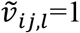 for all *l*. This is plotted in Figure 2E. This has important implications on experimental design for further breeding to explore heterosis.

If we would use this panel of inbred lines as a breeding population, based on the inferred genetic models for different traits we can explore computational means to optimize breeding experimental designs. In maize breeding, the selection scheme is reciprocal recurrent selection (RRS), *i*.*e*., repeated crosses within each heterotic group to create new recombinant inbred lines (RILs) and tested with candidates of the opposite heterotic group for improving breeding gains. This is aimed to preserve and enhance the heterosis while improving inbred line performance as well. Given a comprehensive estimation of QTL positions (*τ*_*k*_, hence *r*_*kl*_ estimated recombination frequencies between QTL), genotypes 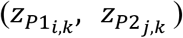 and effects (*a*_*k*_, *d*_*k*_, *aa*_*kl*_) for all inbred lines and hybrids for the targeted traits, we can compute the consequence (in probability) of a cross between any two lines within heterotic groups for line development (genome reshuffling to develop new RILs) or between heterotic groups for choosing parents (testing hybrid). Hence this would be a very efficient computational searching strategy for the selection of mating pairs, a big advantage. This can be combined in a targeted simulation study to explore the short and long-term breeding strategy in this population. This topic will be explored elsewhere.

### Comparison of heterosis in maize and rice

Hua et al. (2002, 2003) reported a rice heterosis study. Starting with a cross between Zhenshan 97 and Minghui 63, the two parental inbred lines that produced F_1_ Shanyou 63, the most widely cultivated hybrid at the time, 240 F9 RILs were produced by single seed descent. RILs were then randomly paired to produce 360 crosses, called immortalized F2 (IMF2). Originally 231 molecular markers were genotyped in RILs and later with extensive genome sequences an ultrahigh-density marker coverage was obtained to produce 1619 bins for RILs (Zhou et al. 2012). Both RILs and IMF2 were planted together and measured for four traits (yield, tiller, grains, and grain weight).

For this population, we orient marker genotypes of P_1_ (Zhenshan 97) as *A*_*l*1_*A*_*l*1_ and those of P_2_ (Minghui 63) as *A*_*l*2_*A*_*l*2_ for all *l* markers. After removing uninformative data, 209 RILs and 276 IMF2 as well as P_1_ (Zhenshan 97), P_2_ (Minghui 63) and F_1_ (Shanyou 63) with a total of 488 sample were used for genetic model fitting and estimation in a joint analysis. Thus, let *s* ∈ (*P*_1_, *P*_2_, *F*_1_, RILs and IMF2). The genetic model for a quantitative trait *y*_*s*_ is defined as

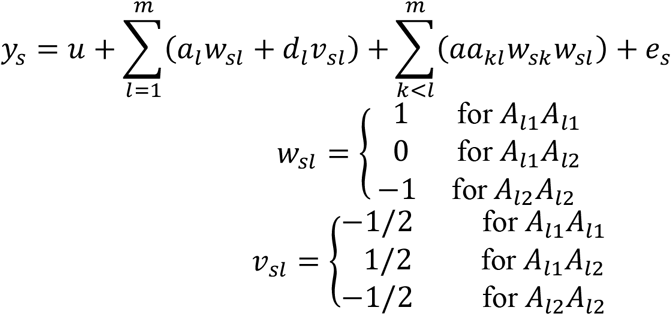

This is the F2 genetic model (G2A model with *p*_*k*_ = 1/2) as the population is an F2-type segregating population. The heterosis is 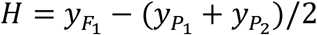, and is expected to be 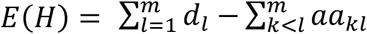 (Melchinger et al 2007, Garcia et al 2008). Also 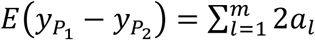. Thus, by estimating the genetic model parameters (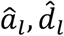 and 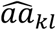) we can compare the estimated and observed heterosis and the parental difference on a quantitative trait.

For this dataset, we used the same model selection procedure as for the maize dataset, except that 80% (rather than 50%) of the samples without replacement were drawn for stability selection. This is to ensure enough QTL with epistasis selected for the final estimation for a proper interpretation of the genetic basis of heterosis, as the sample size in the rice dataset is much smaller than that in the maize dataset.

The selected genetic model information is in supplemental Table S2. The main results are in Table 2 which shows the comparison of the observed and estimated heterosis and parental mean differences for the four traits. Table 2 shows that they agree with each other very well, a general property of the model-based estimation. For yield, QTL dominance contributes 98% of heterosis and epistasis contributes 2% of heterosis. However, for grains, the contributions of QTL dominance and epistasis are similar (48% *vs*. 52%). Table 2 also reports the estimates of the mean degree of dominance 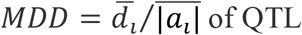 for the four traits.

Figure 4 shows the selection target of QTL effects (*T*_*a*_, *T*_*d*_, and *T*_*aa*_) for the four traits. However, it needs to be pointed out that due to the relatively small sample size, the model fitting and estimation for the rice data are not stable. The sample size of this study may be insufficient to have a reliable model-based inference with epistasis. If we remove QTL epistasis, an estimation of QTL additive and dominant effects is stable.

**Figure 4:**
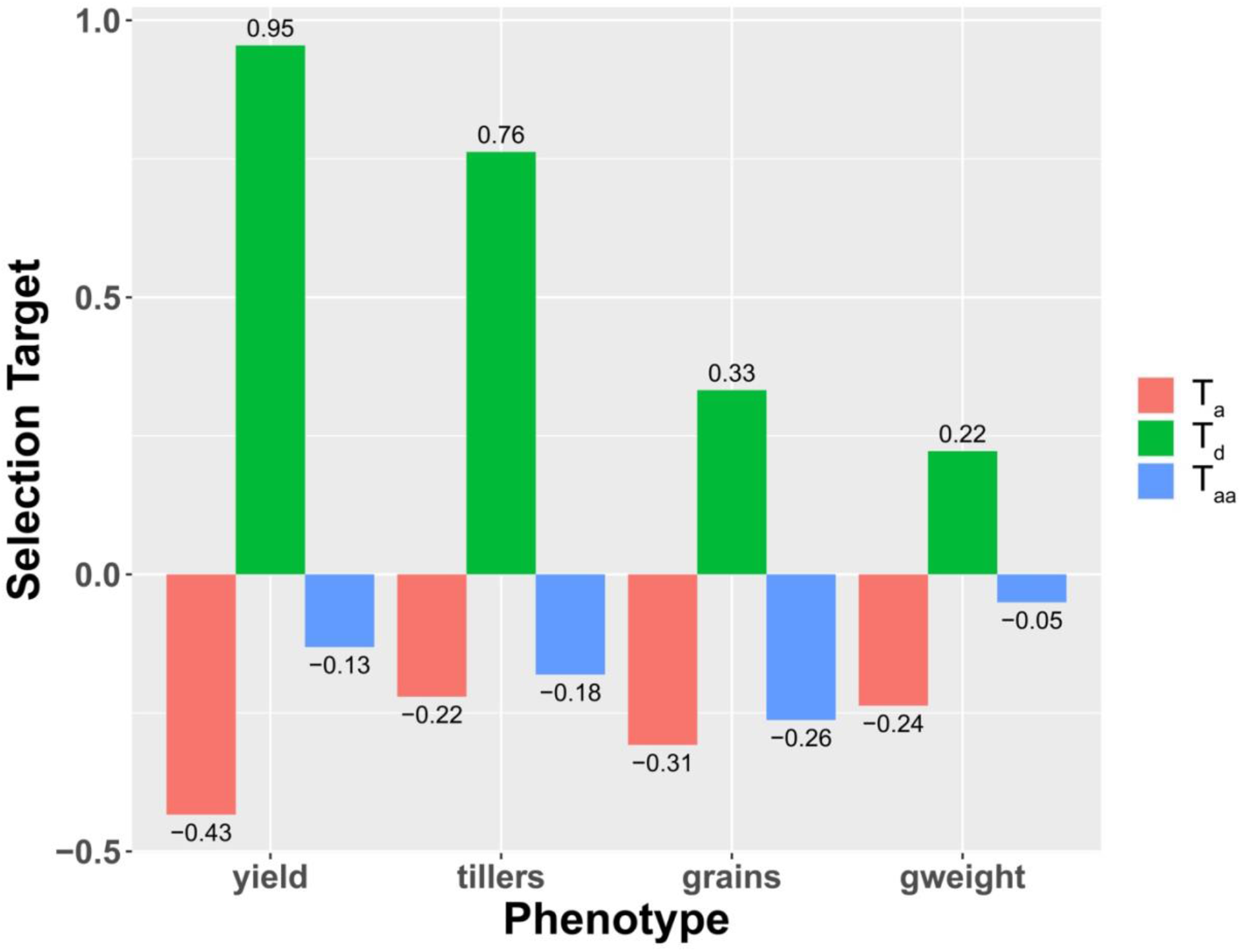
Selection target of QTL effects (*T*_*a*_, *T*_*d*_, and *T*_*aa*_) for the rice study.

### Advantages of G2A model

A genetic model comes with a genetic explanation and thus can be used to explain many genetic phenomena, such as heterosis, as demonstrated in this study. This is the main advantage of a model-based genetic inference. This G2A model can fit to any population and thus can be conveniently used to study the genetic structure of a population. As the G2A model is a binary model, the genetic structure can be represented through a bifurcating tree structure with tree branches proportional to genetic effects and interaction networks connecting tree leaves. Combined with external variables in time (*e*.*g*., pedigree, breeding lines, response to selection) and space (*e*.*g*., environments), this genetic structure can help to illuminate the evolutionary process and to project the future.

## Conclusion

We presented a general theory to analyze and interpret the genetics of heterosis. We applied it to the dataset of a maize study that is a factorial design between a group of male and female inbred lines and their hybrid crosses, and to a rice study that is an F2-type population derived from a cross between two inbred lines and the F1 hybrid. The conclusion on the relative contribution of dominance *vs*. epistasis to yield heterosis is clear. For maize ear weight, the main contribution is QTL dominance (over-dominance to be precise) and the contribution of epistasis is relatively minor. For rice yield, the main contribution is still QTL dominance.

What we presented in this paper is a vision for general quantitative genetic data analysis and interpretation. First, we need to recognize that the fundamental genetic basis of quantitative trait variation is multiple genes. Thus, only when those QTL were identified and fitted in a model for a joint estimation and interpretation, can we have a fuller understanding of the genetic structure in a population, particularly pertaining to the history and evolutionary process that brought about the population. In this inference, a quantitative genetic model or QTL model is the key and the bridge that connects genome to phenome. This connection can implicate the past—evolutionary history. It is the genome that connects individuals, populations, and past, and it is phenotypes that are directly subject to (natural or artificial) selection or other evolutionary driving forces. The inference of the genetic structure in a population, represented by an inferred QTL model, can also be used to project the future that can be explored for a more efficient breeding design and selection scheme in the context of plant and animal breeding.

The underpinning of this vision is that QTL are the elements of quantitative trait variation. The mapping of QTL establishes the physical link to the underlying genes and the joint estimation of QTL effects can reveal the properties of gene actions in a population. This joint estimation and inference are broadly speaking statistically consistent and evolutionary continuous and thus is the right path for general quantitative genetic analysis and interpretation.

## Data availability

The original experimental data were from Xiao et al. (2021) and Zhou et al. (2012). The information of the inferred genetic model is provided in supplemental Table S1 for the maize study and supplemental Table S2 for the rice study and is the basis for all the Figures and Tables. The R codes of the analysis is available upon request.

## Acknowledgments

ZBZ conceived the idea and wrote the paper, GSG and LJM performed the data analysis, YJX and JBY provided the data and stimulating discussion that led to this study. All contributed to the revision of the paper. We thank the reviewers for the constructive comments.

